# Similar time course of fast familiarity and slow recollection processes for recognition memory in humans and macaques

**DOI:** 10.1101/2020.01.10.901975

**Authors:** Zhemeng Wu, Martina Kavanova, Lydia Hickman, Fiona Lin, Mark J. Buckley

## Abstract

According to dual-process theory, recognition memory performance draws upon two processes, familiarity and recollection. The relative contribution to recognition memory are commonly distinguished in humans by analyzing receiver-operating-characteristics (ROC) curves; analogous methods are more complex and very rare in animals but fast familiarity and slow recollective-like processes (FF/SR) have been detected in non-human primates (NHPs) based on analyzing recognition error response time profiles. The relative utility of these methods to investigate familiarity and recollection/recollection-like processes across species is uncertain; indeed, even how comparable the FF/SR measures are across humans and NHPs remains unclear. Therefore in this study a broadly similar recognition memory task was exploited in both humans and NHPs to investigate the time course of the two recognition processes. We first show that the FF/SR dissociation exists in this task in human participants and then we demonstrate a similar profile in NHPs which suggests that FF/SR processes are comparable across species. We then verified, using ROC-derived indices for each time-bin in the FF/SR profile, that the ROC and FF/DR measures are related. Hence we argue that the FF/SR approach, procedurally easier in animals, can be used as a decent proxy to investigate these two recognition processes in future animal studies, important given that scant data exists as to the neural basis underlying recollection yet many of the most informative techniques primarily exist in animal models.

## Introduction

Recognition memory, a form of declarative memory according to some authors (Squire and Zola-Morgan 1991), allows us to make judgements as to the prior occurrence of stimuli based on previous encounters. The delayed matching-to-sample (DMS) and delayed nonmatching-to-sample (DNMS) tasks of recognition memory have been used extensively in animal models. Both tasks are initiated by requiring animals to view the sample picture (or as occurred in many earlier studies, by presenting real objects briefly to animals in a Wisconsin General Test Apparatus), and then after a delay in which no stimuli are presented on the screen, typically two choice pictures are presented in which animals should select the stimulus seen earlier as sample (i.e. the ‘match’) in DMS task while animals should choose the picture different from the sample (i.e. the ‘non-match’) in DNMS task. By manipulating the length of delay between sample and test images, which is often varied within an experiment to test for delay-dependent deficits that might index memory processes; and/or by increasing the number of items to be remembered, the difficulty and cognitive demands on both tasks increase. As in both tasks, studied images will be represented in the test phase, animals may make use of familiarity to differentiate studied and unstudied images. As familiarity is often considered as an automatic process (Jacoby 1991), animals may just rely on familiarity of studies images to solve recognition tests. Whereas animals can clearly judge relative familiarity of stimuli, the issue of whether animals can ‘recollect’ discrimination per se has been a more contentious issue.

Dual-process signal detection (DPSD) theory suggests recognition memory may draw upon two processes, commonly referred to as familiarity and recollection (Dede et al. 2014; Parks and Yonelinas 2007; Scalici et al. 2017; Wixted 2007; Yonelinas 2001, 2002; Yonelinas and Parks 2007). In recollection, if the item/object is judged as being encountered previously, the related contextual information about the item/object can be recalled with more details. For instance, recollection of an item/object may prompt retrieval of information about when and where one was when one encountered the item/object, what one was doing, thinking or feeling, as well as the surrounding environments. By contrast, familiarity is simply a feeling of the item/object has been encountered and generates a feeling of familiarity or knowing but without too much details (Dede et al. 2014; Parks and Yonelinas 2007; Scalici et al. 2017; Wixted 2007; Yonelinas 2001, 2002; Yonelinas and Parks 2007). Accordingly, recollection versus familiarity is often characterized as remembering versus knowing. The receiver-operating-characteristics (ROC) analysis in humans has shown two different components in human recognition memory: a symmetrical and curvilinear component associated with familiarity, and an asymmetrical and linear component related to recollection (Parks and Yonelinas 2007; Eichenbaum et al. 2007; Yonelinas and Jacoby 1994; Yonelinas 2001, 2002; Yonelinas and Parks 2007; Yonelinas et al. 2010).

Recent animal studies have also investigated familiarity and recollection-like memory processes by analyzing the ROC curves based on dual-process signal-detection (DPSD) model (Fortin et al. 2004; Guderian et al. 2011; Sauvage et al. 2008). A fundamental difficulty to overcome in animal models using ROC is that one cannot simply ask for a confidence judgment after the recognition memory judgment. The first application of ROC curves to distinguish familiarity and recollection-like processes in animals was in the rodent model wherein Fortin et al. (2004) plotted ROC in rodents by varying the reward and effort required to make old versus new judgments so to allow responses to be assessed across different levels of ‘confidence’. Fortin et al. (2004) showed that these ROC looked like ROC in normal humans; moreover, in rodents with hippocampal lesions the changed shape of the ROC resembled the changed shape of ROC in human patients with hippocampal lesions (i.e. asymmetrical curvilinear ROC became curved symmetrical ROC). In patients this change may be interpreted as loss of recollection but spared familiarity; hence in rodents with hippocampal lesions Fortin et al (2004) showed that they too possibly lost a recollective-like component to their memory but their familiarity remained intact (Fortin et al. 2004). Subsequently, Sauvage et al. (2008) used an associative recognition paradigm and ROC to show recollection is reduced and familiarity increased after hippocampal lesions in rats (Sauvage et al. 2008). These issues are not without controversy and opposing views generated lively debate (Eichenbaum et al. 2005, 2008; Wixted and Squire 2008). Later, Guderian et al. (2011), by biasing macaques to respond to new and old stimuli by altering reward amounts they could obtain for their correct responses, further confirmed, using ROC, that recognition memory in animals draws upon separate familiarity and recollection-like processes in macaques (Guderian et al. 2011).

A key alternative approach to investigating the relative contributions of familiarity and recollection have investigated the time course of familiarity and recollection processes. Time courses of recollection and familiarity have also largely been investigated in human studies. For example, Yonelinas and Jacoby (1994) calculated the peak time of familiarity and recollection in a source memory task and showed that whilst the peak time of familiarity is faster, between 600 and 800 ms, the peak time of recollection is slower, between 800 and 1000 ms; this gives rise to the concept of faster familiarity and slower recollection (Yonelinas and Jacoby 1994). A faster response to retrieving familiar words (i.e. familiarity process) has likewise been found in comparison to retrieving details of words in a word list (i.e. recollection process) (Hintzman and Curran 1994). Some neurological studies have also confirmed an earlier process of familiarity compared to a later process of recollection in recognition memory. For instance, Curran (2000) found the components of event-related brain potentials (ERPs) related to familiarity appears earlier at 300-400 ms than other later parietal components associated with recollection at 400-800 ms (Curran 2000).

Dosher and his colleagues applied a speed-accuracy trade-off paradigm to interpret the time course of these two processes, in which human participants must respond within a brief window following a go-signal, which is presented at various delays after the offset of the encoded sample items (Dosher and Rosedale 1991; Dosher 1984). In this paradigm, a plot of false alarm errors against retrieval time shows a rise of errors across extremely short retrieval time that reaches a summit in the intermediate retrieval time then decreases in the longer retrieval times (Brainerd and Reyna 2005; Dosher 1984; Dosher and Rosedale 1991; Gronlund and Ratcliff 1989; Hintzman and Curran 1994; Matzen et al. 2011; Rotello and Heit 2000; Rotello et al. 2000). The peak time of these false alarm errors has been considered to relate to the onset of familiarity, and the suppression of false alarm errors during the longer moderate/intermediate retrieval times is considered to reflect the onset of recollection which is conducive to minimizing false alarms (Brainerd and Reyna 2005). Taken together, there is significant consensus as to “fast familiarity” and “slow recollection” (FF/SR) across the human literature, although the methodology of speed-accuracy trade-off paradigm has been questioned in some other human studies (Brainerd et al. 2014, 2019).

More recently, Basile and Hampton (2013) have reported similar FF/SR profiles in non-human primates (NHPs). Their study is similar to the speed-accuracy trade-off paradigm used in human studies (Dosher 1984; Dosher and Rosedale 1991), wherein the time course of familiarity and recollection are investigated by plotting false alarm errors against retrieval time. Basile and Hampton (2013) introduced a modified recognition memory paradigm for NHPs in which animals were trained to select/touch a standard non-match button on the touch-screen if they wished to indicate that the test image did not match the studied image, but were trained to select/touch the test image itself if they wished to indicate that the test item was a match to the studied image. By this approach this paradigm allowed miss and false-alarm errors to be distinguished which better facilitates application of signal detection theory to the NHP DMS task. By plotting false alarm errors against animals’ response time, the authors found that recognition choices in the recognition memory task could be categorized in three different ways depending on response time: (i) short-latency errors induced by false alarms to familiar lures, deemed to indicate the process of familiarity; (ii) medium-latency responses are less likely to be affected by false alarms and are more accurate, suggesting the onset of recollection-like process that could correctly reject familiar lures; and (iii) long-latency and low accuracy responses which are deemed to be guesses (Basile and Hampton 2013).

There have been criticisms of applying the DPSD model to differentiate familiarity and recollection-like processes in animals, Firstly, the DPSD model assumes that recollection is a threshold process with high confidence judgments in which human subjects either recollect or do not recollect a stimulus; while familiarity is a signal-detection process with various confidence judgments. As whether animals can ‘recollect’ discriminanda per se is still debated as discussed above, though at least some studies are favorable to this view. Secondly, animal models have had to develop proxies for humans’ reported confidence judgments. As outlined above, rats were biased to respond to new and old odour by altering the amounts of rewards in different heights of test cups (Fortin et al. 2004; Sauvage et al. 2008); and macaques were biased to respond to new and old stimuli by altering different amounts of rewards they could obtain for their correct responses (Guderian et al. 2011). In the above three animal studies, it is assumed that animals have to be sufficiently confident to overcome any given level of bias, so the bias levels can be used to estimate their confidence. However, training animals to respond to correct new and old stimuli based on such bias levels so as to generate ROC is difficult and time-consuming and we are not aware of other studies that have achieved this to-date; whilst an important and influential methodology it is unlikely to become a frequently used methodology in animal models.

Other criticisms on interpreting animal recognition memory using DPSD models are focused upon the model itself. Malmberg (2008) points out the lack of consistency in the measurements of recollection and familiarity derived from DPSD model even across different human studies (Malmberg 2008). A review from Yonelinas and Parks (2007) raised several issues related to this inconsistency which should be carefully considered when applying DPSD to animal models. A first considers the average performance rate; either performing too well or too poorly will result in the ceiling and floor effects, which will lead to the difficulties in ROC curve fitting in the DPSD model and makes the interpretation of familiarity index based on slope of ROC less certain (Yonelinas and Parks 2007). In animal studies, it may be harder to control the overall performance of the animals since most animals are over-trained in behavioural tasks. Secondly, as with human participants, if the entire scale of confidence judgments is not utilised, then ROC curve fitting is less reliable and parameter estimates extracted from ROC curve will be less meaningful (Yonelinas and Parks 2007). Moreover, in animal models, if animals showed significant preferences to trials with certain biases values (i.e. trials with higher biases values representing larger rewards in studies of Fortin et al., 2004 and Guderian et al., 2011 then the number of reliable data points to fit ROC curves may decrease. Additionally, curve fitting results are easily influenced when hits or false alarm rates on the extreme ends of the ROC approach 0.0 or 1.0 (Yonelinas and Parks 2007); negative values of R (recollection index) or F (familiarity index) may be reported but are meaningless considering the nature of recollection and familiarity process. All the above issues are challenges for DPSD model-based interpretation of human ROC and have arguably larger impact upon animal ROC studies of recognition memory.

The FF/SR paradigm on the other hand is relatively easy to implement in both humans and NHPs (Dosher 1984; Dosher and Rosedale 1991; Basile and Hampton 2013). In the NHP version of this paradigm reviewed earlier (Basile and Hampton 2013) only two images were used repeatedly, in each trial, so in each trial one was the studied image to remember while the other one the lure (or vice versa). Basile and Hampton (2013) argued that this was appropriate larger sets are easier to remember (i.e. to avoid too few errors for analysis) and also that large sets may be too easily discriminabale on basis of familiarity; however this was not verified in that study. Certainly small sets are likely to elicit greater proactive interference but this is expected to be very limiting indeed for recollection and familiarity per se; the task essentially becomes a recency memory task which weakens the interpretation of Basil and Hampton’s data. Therefore our study of FF/SR here expands upon Basil and Hampton (2013) by looking at both small sets and larger sets (trial-unique) of stimuli, to overcome the aforementioned limitation and more importantly, seek generalization of FF/SR effects across a range of set-sizes.

In this study, we adopted Basile and Hampton’s basic recognition memory paradigm, but further refined the paradigm to probe intra-species comparisons across humans and NHPs. Firstly, in human subjects, we used the tasks to assess whether response time derived measurements of recognition errors could be used to differentiate fast familiarity and slow recollection processes in humans using similar paradigms to that we used in NHPs. Then, to probe the relationship between recognition error response time (FF/RS) derived and ROC derived estimates of familiarity and recollection indices in humans, we first sorted trials from slowest to fastest in order to assign trials to a series of response time bins (ranging from fast to slow), and then fitted human ROC data to the DPSD model and extracted familiarity and recollection index independently across the different response time bins. To enable comparison of approaches we then plotted recognition errors (i.e. miss errors and false alarm errors) independently across similar response time bins to those used in the ROC approach. We compared the plots of recognition errors to plots of ROC derived familiarity/recollection indices along a common time course. Following Basile and Hampton (2013) we hypothesized a peak of false alarm errors associated with the short latency bins and we further hypothesized this ‘fast-familiarity’ behaviour would correspond with a peak of ROC-derived familiarity index; accordingly we also hypothesized that a peak in recognition accuracy (d prime) would correspond in time-course with the peak of the ROC-derived recollection index in the medium-latency bins. If the aforementioned hypotheses were supported then, this would suggest that the time course analyses of patterns of recognition errors are phenomenologically related to the ROC-derived familiarity and recollection indices. Then, benefiting from using a similar recognition memory paradigm in both humans and macaques (Fig.1), we were able to compare plots of recognition errors against response time in humans to those in NHPs on a similar task so to investigate the time course of familiarity and recollection across species; accordingly we hypothesized similar “fast familiarity” (peak of false alarm errors) and “slow recollection” (peak of accuracy) profiles across response time in both species. If our first hypothesis about a relationship between recognition error-derived and ROC-derived familiarity/recollection indices holds then this significantly increases the validity of the response time approach derived from recognition errors in NHPs for probing their familiarity and recollective-like (i.e. dual-process) contributions to recognition (of clear benefit to future behavioural neuroscience research in animal as to the neural basis of recollection given the inherent difficulties associated with fully exploiting ROC approaches in animals as we reviewed earlier).

**Figure 1.**
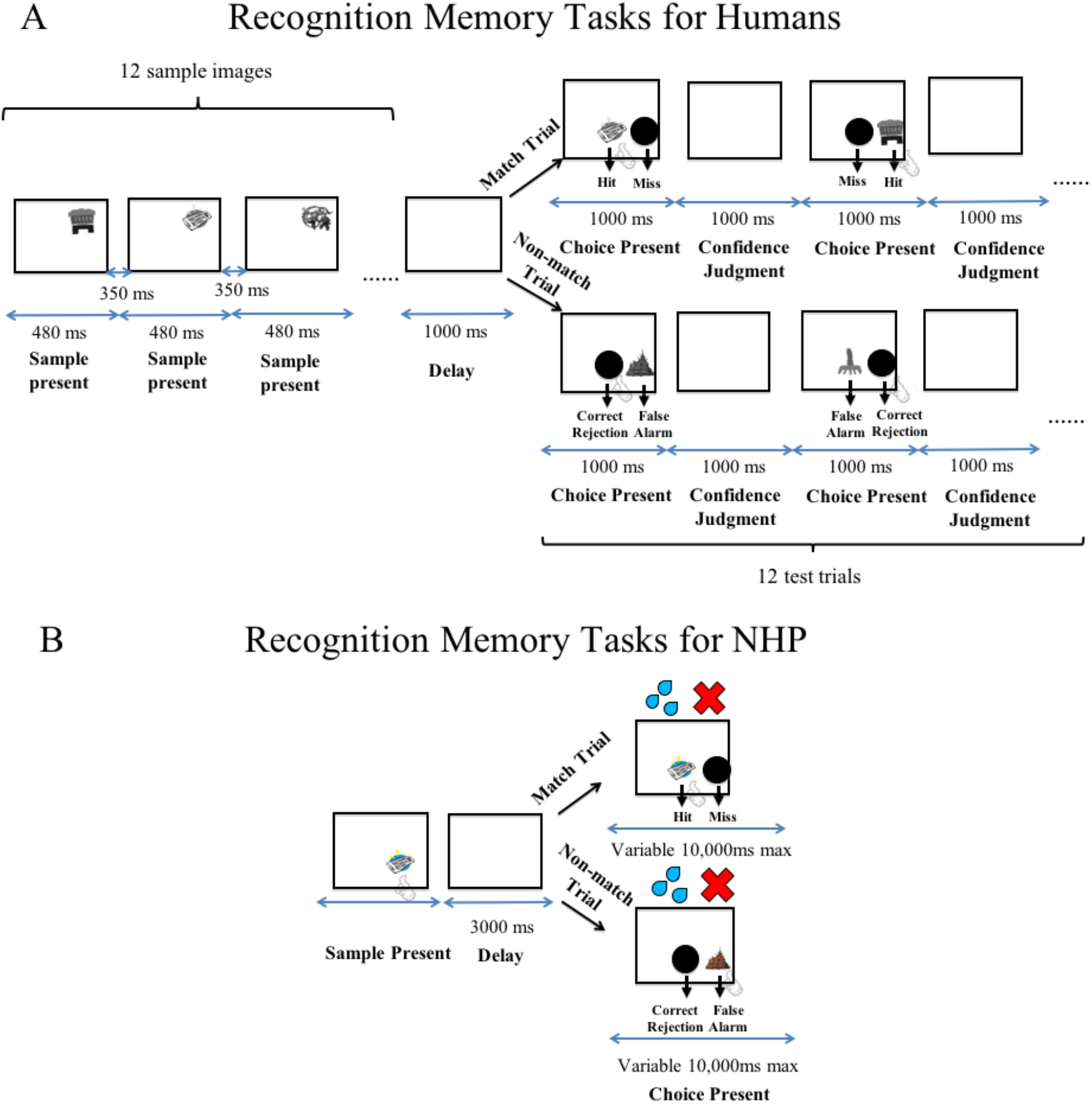
Diagram of the match/non-match recognition memory paradigms for human participants (top panel, A) and NHPs (bottom panel, B). A, in the recognition memory tasks for humans, participants were instructed to remember 12 gray-scale sample images, then after a delay of 1000 ms, they were instructed to touch the test image test image if it matched the sample image; or selected the standard ‘non-match button’ if not match; and then report their confidence judgments using hand gestures in each test trial; they completed blocks of 12 sample trials and then 12 test-trials in this way (15 blocks in total per session). B, in the NHP recognition memory tasks the NHP viewed one sample image, and experienced a delay of 3000 ms, and then was trained to touch the test image if it matched or to select a standard non-match button if not match; this trial structure repeated throughout the session with some samples and test-items being novel (trial-unique) and others familiar within (but not between) daily sessions.

## Results

### Experiment 1: Human behavioural study

Performance data from all 27 participants were included in the analysis; each participant completed 180 trials (12 test trials in each of 15 blocks). For analysis of recognition errors, for each participant, we first grouped their 180 trials into 10 response time (RT) bins (therefore each bin contained 10% of each participant’s trials), and then calculated false alarm error rate, miss error rate, and d prime (an index of discriminability/accuracy) independently for each RT bin.

D prime (d’) was calculated according to the output of an equal-variance signal-detection (EVSD) model wherein new-items and old-items are represented by two Gaussian distributions along the dimension of memory strength with equal variance. The d’ measure depends both on the separation (difference in standardized means of new-item and old-item distributions) and the spread (standard deviation) of new-item and old-item distributions. As the EVSD model assumes the variance of new-item and old-item distributions are equal, d’ is effectively the distance between new-item and old-item distributions (expressed as a z score with respect to its magnitude relative to the standard deviations of the distributions), reflecting sensitivity or discriminative ability to judging old-items from new-items. In our recognition error analysis, d′ was calculated in each RT bin, as an indication of discriminability/accuracy (see Table 1 EVSD model for mean RTs represented in each bin).

**Table 1.** Mean response latencies for binned human data in Fig.2. Each bin contained 10% of the responses of each participant. For each response bin, we calculated the mean response latency in milliseconds for each human participant, and then averaged those mean latencies to produce a group mean latency for each bin.

| **Bin Number** | **1** | **2** | **3** | **4** | **5** | **6** | **7** | **8** | **9** | **10** |
| --- | --- | --- | --- | --- | --- | --- | --- | --- | --- | --- |
| **EVSD model** | 1074.7 | 1225.0 | 1337.2 | 1443.4 | 1559.3 | 1685.0 | 1834.8 | 2050.8 | 2358.2 | 3089.1 |
| **DPSD model** | 1048.5 | 1209.4 | 1327.4 | 1442.9 | 1554.5 | 1677.0 | 1839.6 | 2067.0 | 2358.0 | 3058.7 |

**Figure 2.**
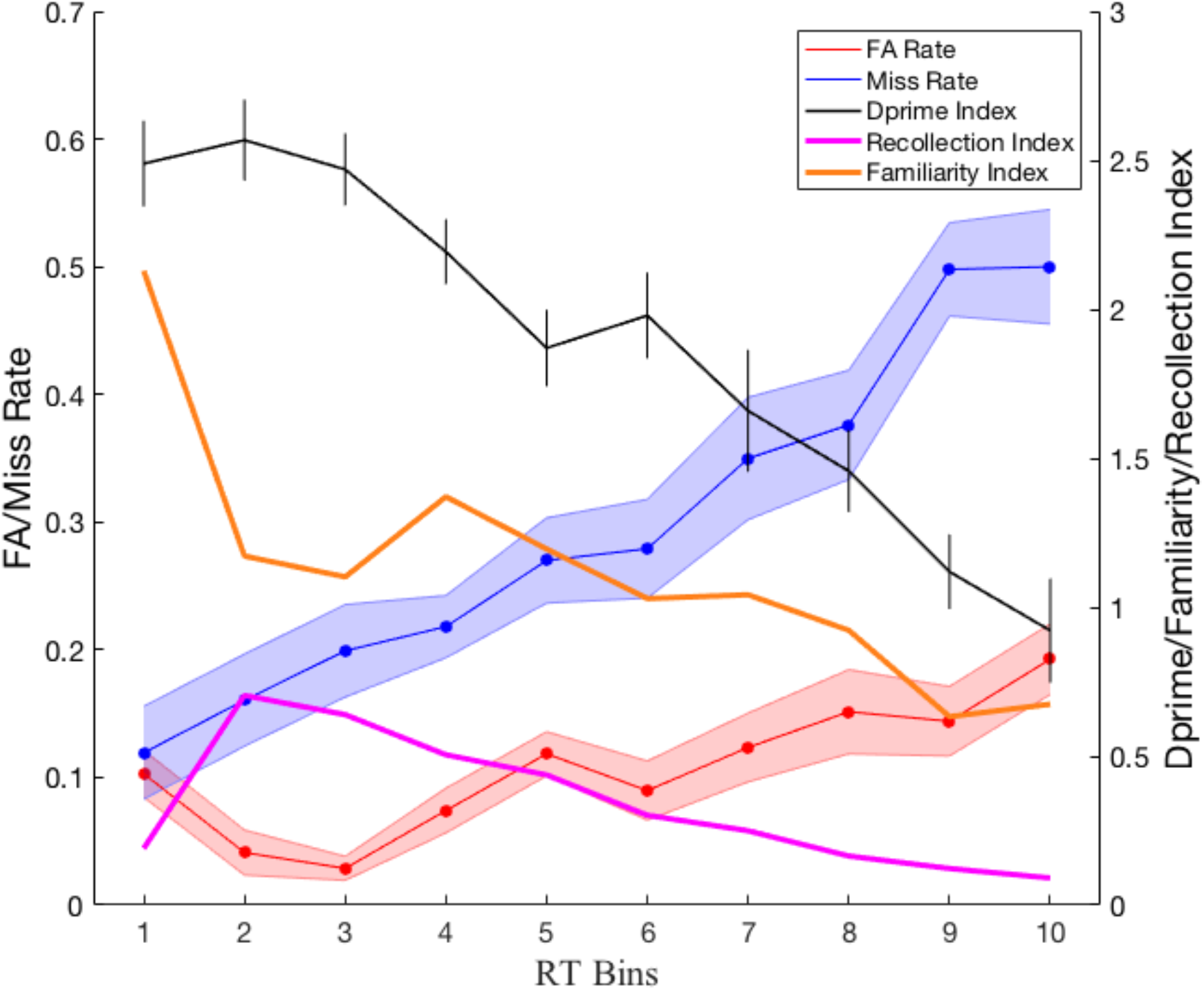
Recognition errors, discriminability/accuracy d prime, and familiarity/recollection indices as a function of response time in humans. Each bin contains 10% of each participant’s trials (ranging from bin 1 which contains the fastest 10% of trials to bin 10 which contains the slowest 10% of trials). Error bars are ± 1 SEM. False alarm errors (red solid line) and miss errors (blue solid line) were almost the same in the first RT bin; then miss errors increased, while false alarm errors initially decreased and then later increased across RT bins. Based on EVSD model, accuracy (d prime, black solid line) demonstrated a slight inverted U-shape across RT bins and reached the summit in the second RT bin. Based on DPSD model, recollection index (magenta solid line) demonstrated an inverted U-shape against the 10 latency bins; while familiarity index (orange solid line) showed a decreasing trend, whose peak value appeared in the first RT bin.

Similar to plots of recognition errors and d’ found in Basile and Hampton’s macaque study (Basile and Hampton 2013), we found that our human participants (Fig. 2) exhibited a slight U-shaped function: their false alarm error rate was significantly higher in our shortest latency RT bin than in the next two RT bins in which RT was lower (first vs. second bin: paired t-test, t_(26)_ = 2.431, *p*= 0.022; first vs. third bin: paired t-test, t_(26)_ = 3.338, *p* = 0.003), after the third RT bin it rose again (third vs. forth bin: paired t-test, t_(26)_ = −2.656, *p* = 0.013; third vs. tenth bin: paired t-test, t_(26)_ = −5.896, *p* < 0.001). On the other hand their miss error rate increased (left to right on Fig. 2) from the fastest RT bins to the slowest RT bines (first vs. tenth bin: paired t-test, t_(26)_ = −6.214, *p* < 0.001). Discriminability/accuracy, measured as d′, demonstrated a slight inverted U-shape across the 10 latency bins, whose peak-to-valley value was significantly larger than zero (second vs. tenth bin: paired t-test, t_(26)_ = 6.958, *p* < 0.001). It should be noted that the peak value of d’ (d’ = 2.57) appeared in the second RT bin (average time: 1225.0 ms). The peak time of false alarm error occurrence has been considered to correspond to the onset of familiarity (Basile and Hampton 2013), and the peak time of discriminability/accuracy (in parallel with a suppression of false alarm rate due to recollection/recollective-like memory processes) was considered to evidence the onset of recollection (Basile and Hampton 2013); accordingly, the patterns and time profiles of recognition errors against response time in humans were similar to the ones found in the previous macaque study of Basile and Hampton (2013) which also showed evidence for fast-familiarity and slow-recollection (FF/SR) in a similar behavioural task.

To further investigate the latency of recollection and familiarity in human participants, we looked for correspondence between the aforementioned analyses and the ROC-derived approach for deriving familiarity and recollection indices. So we next applied the DPSD model to ROC data derived from each of the ten RT bins separately. The DPSD model is more complex form of model containing a single threshold process of recollection and a pure signal-detection process of familiarity; below is the algorithm of this model (Koen et al. 2017) we used:

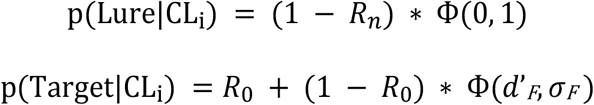

Briefly, in the target and lure distributions (Ф is the cumulative response function), *R_0_* is a target threshold parameter labeled recollection of old stimuli, and *R_n_* is a lure threshold parameter labeled ‘recollection’ of lures. As recollection is assumed to be a threshold process in the DPSD model, stimuli with a strength value above the *R_0_* threshold will be classified as a target. In the DPSD model, as we don’t care about the lure distribution we make it a standard normal Gaussian function with a mean of 0 and a unit standard deviation (*R_n_* = 0). The other two parameters, *d’_F_* and *σ_F_*, are estimates of familiarity. In the DPSD model we used, familiarity is defined as a signal-detection process in which memory strength of target and lure items falls into Gaussian functions. In our DPSD model, we assume variances of target and lure items are equal (*σ_F_* = 1), thus familiarity is defined as an EVSD model with its parameter *d’_F_*. Based on the above assumptions, the algorithm of DPSD model becomes:

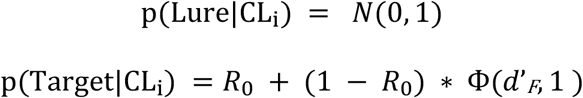

When fitting ROC data to the DPSD model, sufficient trials are required to make ROC curve fitting sufficiently reliable (Yonelinas 2002; Yonelinas and Parks 2007). Therefore, for each human participant, we took the 180 trials divided into 10 RT bins from fastest to slowest and to ensure sufficient trials in each RT bin for curve fitting, trials in the corresponding RT bins of different participants were all combined. So, for example, the 10% fastest trials from each participant came to bin 1, and 10% slowest trials from each participant came to bin 10. Therefore, for analysis, there were 486 trials (i.e. 10% of total number of trials from all 27 participants) in each RT bin. Then for each RT bin independently, a ROC was plotted cumulatively for each confidence level according to the proportion of correct ‘old’ judgements against proportion of incorrect ‘old’ judgements. The recollection index (*R_0_*) and familiarity index (*d’_F_*) were extracted by fitting the model to data minimizing the squared difference between the observed and predicted data in each confidence rating bin. Yonelinas’ research group provides an Excel worksheet which facilitates ROC curve fitting and index extraction (website link: https://yonelinas.faculty.ucdavis.edu/roc-analysis/). In this way, we calculated recollection and familiarity index across 10 RT bins and plotted the values on the same figure as the error-rates (and EVSD derived d′) given both analysis approaches considered 10 RT bins (Fig. 2, see Table 1 DPSD model for average latencies represented by each RT bin).

The recollection index demonstrated an inverted U-shape across the 10 latency bins, whose peak value (recollection index = 0.70) was in the second RT bin (average time: 1209.4 ms), whilst the familiarity index showed a decreasing trend from a peak value (familiarity index = 2.13) in the first RT bin (average time: 1048.5 ms). The mean RTs for recollection/familiarity indexes in each bin is shown in Table 1. It should be noted that the time-course of a significant switch from clearly high familiarity and clearly low recollection to somewhat lower familiarity values accompanied by somewhat higher recollection values appeared from the first to the second RT bin, thereafter both indexes decreased as RT prolonged consistent with the longest latency RT bins being associated with increased rates of ‘guessing’, and also consistent with our observed steady decline in discriminability/accuracy (see plot of d′ on Fig. 2) as RT increases. This is also consistent with the aforementioned evidenced that miss rate steadily increases to 50% across higher RT bins which does indeed corresponds to chance level of performance on match trials. Notably, the average time of peak value of accuracy or d’ in EVSD model was in the second RT bin (average time: 1225.0 ms) and this corresponded to the RT bin (average time: 1209.4 ms) in which the peak recollection index occurred. Taken together our observations provide strong evidence that FF/SR processes exist in humans just as observed by Basile and Hampton in macaques, using broadly similar paradigms. Moreover, the interpretation/conclusions from the FF/SR approach appear consisted with the time profile of indices of recollection and familiarity derived from the ROC approach, further validating the approach.

### Experiment 2: NHP behavioural study

In this experiment, the animal had completed 5 sessions with more than 300 trials per session, and all of the data from these 5 sessions were put into the analyses. Similar to our above analyses on recognition errors in humans, we next analyzed macaque behavioural data across 5 sessions in a similar task; we likewise grouped trials into 10 RT bins, with each bin containing 10% of macaque’s trials. In each RT bin, we calculated false alarm rate, miss rate and d’ (discriminability/accuracy index based on EVSD model). This analysis was done separately for sample images that were novel (Fig. 3B) and those that were familiar (Fig. 3C) to the macaque (also see Table 2 for mean RTs represented in each bin for images as being novel and familiar to the monkey separately).

**Figure 3.**
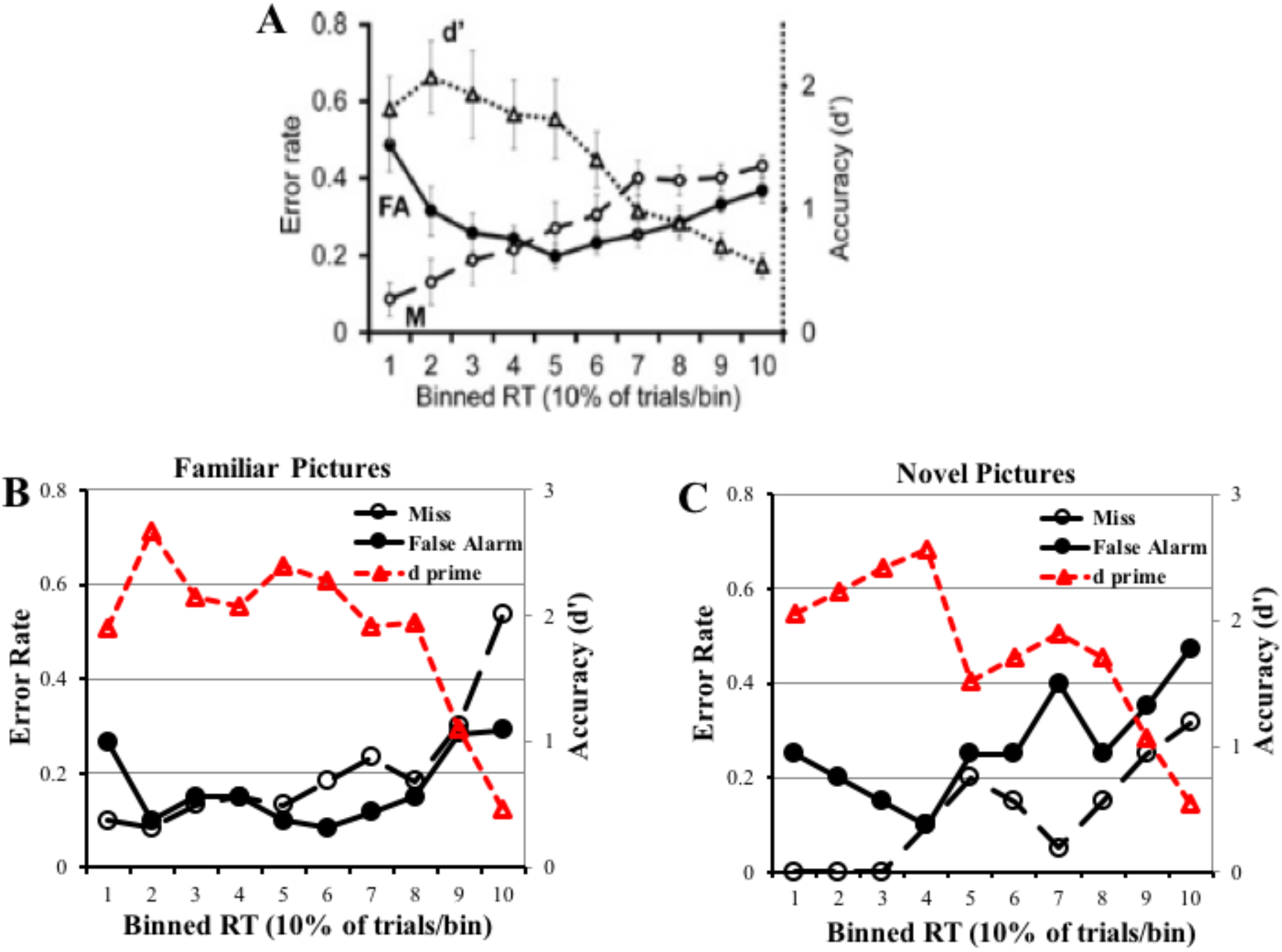
False alarm error rate (solid line with filled circles), miss error rate (dashed line with hollow circles) and discriminability/accuracy d prime (dashed line with hollow triangles) as a function of response time bin in macaques. Each bin contains 10% of each macaque’s trials (ranging from bin 1 which contains the fastest 10% of trials to bin 10 which contains the slowest 10% of trials). Panels A-C depict recognition errors and accuracy from: (A) Basile and Hampton, 2013 study; (B), from our macaque when viewing familiar pictures; and (C) from our macaque when viewing novel pictures. All the panels show that false alarm errors revealed a U-shape function, miss errors increased, and d prime produced an inverted U-shape in relative incidence across the 10 RT latency bins in NHPs.

**Table 2.** Mean response latencies for binned macaque data in Fig. 3. Each bin contained 10% of the responses of the monkey. For each response bin, we calculated the mean response latency in milliseconds for images as being novel and familiar to the monkey.

| Bin Number | 1 | 2 | 3 | 4 | 5 | 6 | 7 | 8 | 9 | 10 |
| --- | --- | --- | --- | --- | --- | --- | --- | --- | --- | --- |
| Novel Image | 781.8 | 936.0 | 1002.2 | 1070.9 | 1180.4 | 1306.2 | 1463.7 | 1657.1 | 1927.8 | 3215.4 |
| Familiar Image | 714.8 | 900.7 | 973.4 | 1048.3 | 1133.5 | 1245.9 | 1384.8 | 1563.8 | 1941.5 | 3689.9 |

Consistent with previous results from Basile and Hampton (2013)’s macaque study which considered only highly familiar stimuli consisting of a repeating small set of just two stimuli (Fig. 3A), our plots of recognition errors for viewing larger numbers of familiar images (Fig. 3B) and also novel images (Fig. 3C) both demonstrated a U-shape function in the false alarm error rate profile across RT bins revealed whereas the miss error rate profile increased across the 10 RT bins.

Discriminability/accuracy, measured as d′ in the EVSD model, produced an inverted U-shape from the fastest RT bin to slowest RT bin, whose peak-to-valley value was larger than zero. Consistent with the main conclusion in Basile and Hampton’s study and with the pattern of recognition errors in our human study (Experiment 1), we found a similar pattern of recognition errors and d′ (discriminability/accuracy) in macaque: at the short response latencies, errors were mostly false alarms consistent with performance being driven primarily by familiarity; at moderate response latencies, the false alarm rate dropped and accuracy was higher consistent with longer-latency recollective-like processes reducing the false alarm rate; and at the longest response latencies, both false alarm rate and miss rate increased and accuracy reached its lowest point consistent with monkeys guessing.

Although the familiarity level of sample pictures didn’t affect the U-shape function of false alarm errors and the inverted U-shape function of d’ as a function of RT, the RT bins in which the minimal values of false alarm errors and the RT bins in which the maximum accuracy (d’) were different for the different familiarity levels. To be specific, the first valley value of false alarm errors (FA rate = 0.1) and the peak value of d’(d’ = 2.66) were in the second RT bin (average time 900.7 ms) for familiar pictures (Fig. 3B); while the first valley value of false alarm errors (FA rate = 0.1) and the peak value of d’(d’ = 2.56) were in the fourth RT bin (average time 1070.9 ms) for novel pictures (Fig. 3C). The mean RTs for novel and familiar stimuli in each RT bin was shown in Table 2. As the peak value of d’ (also corresponding to the first decrease in false alarm error rate) may indicate the start of recollection, it appears that novel images induce a switch from familiarity driven to recollection/recollective-like driven contributions to recognition memory later with respect to time-course, compared with familiar pictures.

## Discussion

This study investigated the time course of familiarity and recollection/recollection-like processes contributing to recognition memory in both humans and NHPs. A first key aim was to examine relationships between ROC derived indices of recollection and familiarity with recognition errors derived measures of ‘fast-familiarity’ and ‘slow-recollection’ (FF/SR) in humans. A second key aim was to compare FF/SR analyses in humans and NHPs. A major motivation for these aims was to strengthen interpretations that may be made from FF/SR studies pertaining to recollective-like behaviour in animals, in existing behavioural studies and future neuroscientific enquiries; this is important because FF/SR data is markedly easier to obtain in animals than ROC data (Fortin et al. 2004; Guderian et al. 2011; Sauvage et al. 2008). In the current study, using a similar task structure across species, human participants and macaques performed an object recognition memory task (with distinct match and non-match trials, see methods, to distinguish two types of errors, false alarms and misses). Our expectations were that: at the shortest response latencies, false alarm errors would be high, attributed to familiarity; intermediate-latency responses would be more accurate and associated with fewer false alarm errors, which is attributed to an onset of recollection or recollection-like processes reducing the false alarm error; at the longest response latencies, both false alarm errors and miss errors would increase in frequency and accuracy would reached its lowest point suggestive of guessing to a greater degree. Prior evidence from human studies suggest that familiarity arises earlier than recollection, for example in studies in which participants used slow recollection to countermand false alarm errors induced by familiarity (Rotello and Heit 2000; Rotello et al. 2000). In our macaque study, most of the short-latency errors were false alarm errors; and at the intermediate response latencies, false alarm rate decreased along with an increase of d’, consistent with a slow recollective-like process to suppress false alarm errors and increase accuracy. In humans, the short-latency errors were occupied by miss and false alarm errors equally; at intermediate response latencies, false alarm errors showed a decreasing trend along with an increasing of d’, which indicates a similar start point of recollection-driven memory. Thus, it is concluded that both humans and macaque can use “recollect to reject” to override false alarm errors driven by familiarity and their response profiles are broadly similar on the FF/SR paradigm.

Robust evidence in support of “fast familiarity and slow recollection” has been provided by many human recognition memory studies (Brainerd and Reyna 2005; Dosher 1984; Dosher and Rosedale 1991; Gronlund and Ratcliff 1989; Hintzman and Curran 1994; Matzen et al. 2011; Rotello and Heit 2000; Rotello et al. 2000). For example, Gronlund and Ratcliff (1989) found little or no evidence for familiarity-based retrieval of studied items at extremely brief retrieval time (i.e. less than 200 ms), but as retrieval time lengthened (i.e. 300-500 ms), familiarity-based evidence accumulated accompanied by increasing false alarm rates, and.as retrieval time became longer (i.e. more than 500 ms), recollection-based evidence was taken into account and accumulated to suppress false alarm rates (Gronlund and Ratcliff 1989; Brainerd and Reyna 2005). Consistent with these studies, our human and macaque results showed a U-shaped curve of false alarm errors against response time as detailed earlier (and see Figs. 2 and 3); however we did not find the corresponding very low false alarm probability at extremely low response times as Gronlund and Ratcliff (1989) and other showed. One possible reason for this might be that our studies didn’t include such extreme short response time periods. Indeed, in the study of Gronlund and Ratcliff (1989) their very short (<600ms) latencies included both sample-to-test item delay plus response time (also see the review by Brainerd and Reyna, 2005) but in our macaque study, the delay between sample and test image was already 3000 ms and in our human study was 1000 ms. Instead, the U-shaped functions in our study, seen also in the macaque study of Basile and Hampton (2013), are consistent with the higher response times (wherein an inverted U-curve in seen) in the study of Gronlund and Ratcliff, in which false alarm errors reached highest point at short response latency and then decreased.

In this study we compared the response time profiles for fast-familiarity (FF) and slow-recollection (SR) in macaques and humans; for each species we plotted recognition errors and discriminability/accuracy (indicated by d’ in EVSD model) against response time (see Figs. 2 and 3). The summit of accuracy for humans appeared in the second RT bin, which indicates that a switch from relying primarily upon familiarity to relying upon recollection has already occurred by around 1225.0 ms (see Table 1 EVSD model for mean RTs represented in the second bin). In Figure 2, to compare ROC-derived indices of recollection and familiarity to the FF/SR interpretations we also plotted curves through the values obtained from deriving familiarity and recollection indices on the data in each bin from fitting ROC data to the DPSD model. The peak values of the ROC-derived familiarity index occurred much earlier in RT Bin 1 (corresponding to average peak time of familiarity index of 1048.5 ms) than the peak values of the ROC-derived recollection index in RT Bin 2 (corresponding to average peak time of recollection index of 1209.4 ms). The timing profiles of the ROC-derived indices of recollection and familiarity correspond with and corroborate the timing profiles of the FF/SR based interpretations further supporting and validating the FF/SR approach in humans. Moreover, the peak ROC-derived recollection index base on DPSD model fell in the second RT bin in which the discriminability/accuracy (d’, calculated based on recognition errors in EVSD model) was also in peak, suggesting recognition error distributions can be used as a proxy for estimating the onsets of recollection processes in humans. In our macaque study, the trends of recognition errors and accuracy were similar to those in humans (compare Figs 3B, C with Fig 2). Accordingly, if the peak time of accuracy can be used as a proxy to estimate the onset of recollection processes within the response time profile, then we estimate that an analogous switch from relying primarily upon familiarity to relying upon recollection in the macaque was at an average time of around 900.7 ms for familiar sample images (Fig. 3B), and with average time of 1070.9 ms for novel sample images.

The extent to which image repetition affects familiarity or recollection is an issue of debate. In some studies with humans repetition of studied items improves both familiarity and recollection; image repetition was found to increase false alarm errors if participants were under pressure to respond quickly but had no effects on false alarm errors if participants had more time to respond (Jacoby et al. 1998; Jacoby 1999; Jones and Jacoby 2001). Other studies using the remember-know (R/K) procedure (Tulving 1985) found that repetition of items enhanced recollection/remembering rather than familiarity/knowing (Mántylá and Cornoldi 2002; Pitarque et al. 2015). In our macaque study, as mentioned above, the estimate switch time from familiarity to recollective-like processes occurred earlier for familiar than novel sample images. It appears that image repetition expedites the onset of recollection-like process so as to enhance the performance of recognition memory in NHPs. In a recent macaque electrophysiological and human ERP study, image familiarization sharpened the response dynamics of inferotemporal neurons of macaques and of human extrastriate visual cortex (Meyer et al. 2014). This finding illustrates that repeated viewing of an image modulated neural responses in both macaque and human visual areas, which may be conducive for efficient coding of stimuli. As we didn’t carry out a similar investigation of image repetition effects on familiarity/recollection in humans, it is worth future studies investigating whether a similar behavior pattern can be found and is so to investigate the neural mechanisms behind the behavior in both species.

A small number of influential animal models (with rodents and more recently macaques) used the ROC approach to assess contributions of familiarity and recollection-like processes to recognition memory. These studies manipulated animals’ biases to respond “old” or “new” to stimuli by varying the relative amount of reward corresponding to correct old and correct new responses across various levels (Fortin et al. 2004; Guderian et al. 2011; Sauvage et al. 2008). As stated earlier, these types of manipulation are complex and time-consuming to train but most importantly the bias levels cannot be treated equivalently to humans’ confidence judgments, with the result that interpretation of ROC curves in animals has been questioned and hotly debated (Eichenbaum et al. 2008; Wixted and Squire 2008). We consider those animal models important and influential but at the same time we recognize both the practical and theoretical limitations. Indeed, this has been a major motivation for the current study which we consider successful in strengthening the interpretation of a proxy for differentiating between familiarity and recollection-like processes in animals, namely by investigating the time-course patterns of recognition errors. Our data further validates this FF/SR approach as providing valuable insight into these processes and hence future investigations of familiarity and recollection-like processes in animals, which are in need of advancing given the scarcity of animal models of recollection/recollective-like memory processes (see also (Basile et al. 2009; Basile and Hampton 2011; Paxton and Hampton 2009; Hampton 2001; Hampton et al. 2004; Hampton and Hampstead 2006) for some other approaches to look at other elements of episodic memory in animals).

In summary, the conclusions from this study are that i) “fast familiarity” and “slow recollection” provide comparable measures in both humans and macaques performing similar tasks; ii) both species applied a similar strategy of “reject to recall” to countermand false alarm errors based on recollection/recollective-like memory processes; and iii) image familiarization expedites switches from familiarity to recollection-like processes in macaque; and iv) the time course of familiarity and recollection can be estimated relative to the time-to-peak value of discriminability/accuracy based on recognition errors, and this estimation method is validated by DPSD models used in humans.

## Materials and Methods

### Experiment 1: Human behavioural study

#### Participants

27 participants (18 male, 9 female, age range 18-30 years) took part in the TMS study. Participants were fluent English speakers, right-handed, and had normal or corrected-to-normal vision. Prior to the study, all the participants provided written consent and went through safety screening check to make sure they had no history of previous or current neurological or psychiatric conditions and were not taking any psychoactive medication. All the participants received monetary compensation for their participation at a standard rate for volunteers in Oxford. This study was carried out with the approval of Medical Sciences Interdivisional Research Ethics Committee, University of Oxford.

#### Task Stimuli and Apparatus

The task was an object recognition memory task similar to the one used in NHPs, similarly programmed using Turbo Pascal (Borland), run under DOS on a desktop PC and presented on a 20.1” colour touchscreen (TFT LCD TS200H GNR). The objects images used in the task were clip-art images as in NHP study, but in order to increase difficulty (in light of a pilot study wherein performance was close to ceiling) the images were all converted to grey-scale and the contrast toned down in an attempt to make them harder to discriminate from each other. Additionally, the samples were presented in lists followed by lists of choice trials, again to reduce ceiling effects in the human version of the NHP task. Each image subtended 10° of visual angle in width and 10° in height to the participant sitting facing the screen. The sample image was always presented on the right top of the screen, positioned +12 ° horizontal and −12 ° vertical from the center of the screen. The test images were presented on the right bottom of the screen: one was positioned 0 ° horizontal and +5 ° vertical from the center of the screen; and the other one was positioned +23 ° horizontal and +5 ° vertical from the center of the screen. The background colour to the screen was white.

Participants sat with their eyes a distance of 25 cm from the screen, wearing earplugs, resting their chins on a chin-rest and their foreheads on a head holder to stabilize their head position throughout the experiment. They were instructed to respond to items by touching them on the screen and gestured their confidence ratings using their right hands. For example, they indicate by raising fingers (1, 2 or 3) whether their opinion corresponding to their being somewhat confident (1), moderately confident (2), or absolutely confident (3) in their judgment as to whether they considered the test-item to be old (i.e. presented before in the preceding list as a sample) or new (not seen before in the preceding list as sample). None of the samples in this task were used in more than one list so all stimuli were trial-unique (and hence compared to the novel/trial-unique stimuli in the NHP task).

#### Behavioural task

Prior to the experiment, participants were given instructions on how to perform the task, and how to indicate their confidence judgments (see above), and then they were introduced to the behavioural task in a short training/practice session (comprised of one list of 12 samples and then 12 test-item trials) of the task without stimulation to provide an opportunity to become familiarized with task procedure. We used lists in the human version to avoid ceiling effects as pilot investigations revealed the NHP task with single samples and single test-trials to be too easy for participants. The experimental session itself contained 3 sub-sessions, with each sub-session containing 15 blocks of trials. Participants took a 10-15 min break between each sub-session. Each block contained an encoding phase (i.e. a list of 12 sample images), and then a short delay, and then a test phase (i.e. a list of 12 test-item trials). The task structure is depicted, for one block, in Figure 1 (top panel, A); during the encoding phase, 12 sample images were presented sequentially on the screen (individually for 480 ms), with inter-stimulus-interval as 350 ms. Participants were instructed to view and try to remember each sample image. After the sample phase (all 12 sample items) was completed, a blank screen was presented for 1 s (delay), and then the test phase commenced (all 12 test trials). Test trials were either ‘match trials’ or ‘non-match trials’. In each test trial, either an identical stimulus to one of the preceding12 samples (i.e. ‘match trials’) was shown as a test image, or a novel and previously unseen image (i.e. ‘non-match trials’) was shown as a test image, and that test image appeared on the screen together with a black circle (the left/right position of the black circle and the test-trial image were randomized between trials). The match and non-match trials were also counter balanced (6 of each per test phase of 12 test-trials) and were put in a random order. Just as in the NHP version of the task (Experiment 2) participants were instructed to touch the test image if they thought it matched one of the 12 sample images, or touch the standard ‘non-match button’ (i.e. the black circle) if they thought the test image was not a match. After responding to the test image, participants were instructed to rate their confidence as to whether the test item was new or old using a scale of 1-3 by making three different movements with their fingers, corresponding to somewhat, moderate, and absolute confidence. Participants were further instructed to try to use the entire range of confidence responses as best they could and not simply select the extremes of confidence as defaults.

### Experiment 2: Macaque behavioural study

#### Subject

One adult female rhesus macaque (*Macaca mulatta, age 8 years, weigh 10-13 kg*) participated in this experiment. All animals in our lab are socially housed (or socially housed for as long as possible if later precluded, for example, by repeated fighting with cage-mates despite multiple regrouping attempts) and all are housed in enriched environments (e.g. swings and ropes and objects, all within large pens with multiple wooden ledges at many levels) with a 12hr light/dark cycle. The NHP always had *ad libitum* water access 7 days/week. Most of its daily food ration of wet mash and fruit and nuts and other treats was delivered in the automated testing/lunch-box at the end of each behavioral session (this provided ‘jack-pot’ motivation for quickly completing successful session performance; supplemented by trial-by-trial rewards for correct choices in the form of drops of smoothie delivered via a sipping tube) and this was supplemented with fruit and foraging mix in the home enclosure. All animal training and experimental procedures were performed in accordance with the guidelines of the UK Animals (Scientific Procedures) Act of 1986, licensed by the UK Home Office, and approved by Oxford’s Committee on Animal Care and Ethical Review.

#### Task stimuli and apparatus

The object recognition memory task was programmed using Turbo Pascal (Borland), run under DOS on a desktop PC. Visual stimuli used in the task were clip-art images in colour, which were presented on a 20.1” colour touch-sensitive screen (TFT LCD TS200H GNR). Those clip-art images used in the task were from a large pool of several thousand unique images. Each image subtended 5° of visual angle in width and 5° in height to the subject when presented on the screen. Both the sample image and test images were presented on the right bottom of the screen. The sample image was positioned +10 ° horizontal and +5 ° vertical from the center of the screen, with equal distance to the two test images. The test images were presented at the same horizontal level as the sample image: one was positioned 0° horizontal and +5 ° vertical from the center of the screen; and the other one was positioned +20 ° horizontal and +5 ° vertical from the center of the screen. The background colour to the screen was white. In each session, images were randomly chosen from the pool without replacement and were not re-used on the other testing days.

The animal was seated in a primate chair (Rogue Research Inc.) in front of the touch screen with its head-fixated and whilst it performed the recognition memory task in a magnetic-shielded, and partially sound-attended, testing-box. A window in the front of the chair provided its access to the touch-screen itself. The distance between the monkey and touch screen was fixed at 50 cm enabling the animal to touch the screen easily. An infrared camera was used to monitor the general status of the monkey in the box. A peristaltic pump device located on top of the box fed smoothie reward through a tube and to a spout positioned in the vicinity of the animal’s mouth. Below the screen was also an automated lunch-box which contained the majority of the animal’s daily meal (wet mash and fruits and nuts etc.) and which opened immediately at the end of the task.

#### Behavioural task

In the NHP study, a similar task design has been exploited with the main difference being samples and test phases were not blocked. In each trial, animal viewed a sample image (shown in the center of the screen) to remember and touched it to progress. Then after the 3000 ms delay period, the NHP was tested with two choice stimuli (one an object image and the other a black circle, left-right randomized between trials). The NHP was required to make a choice to the touchscreen to either the object test-image stimulus or to the black circle stimulus. The object stimulus was either the identical stimulus to the sample seen earlier in the trial or it was not identical to the sample. The animal was rewarded by delivery of 10 ml of smoothie for touching the test-item image if it matched the sample image (these we refer to as ‘match trials’), or it was rewarded for selecting the standard ‘non-match button’ (i.e. the black circle) if the test-item was a non-match (these we refer to an ‘non-match trials’). The next trial started after a 3000 ms inter-trial interval after a correct response, or after a 10 s ‘time-out’ after any kind of error response (or after any failure to hold keytouch for the required time, or after failure to release when required). A schematic of the recognition memory task is illustrated in Figure 1 (bottom panel, B). Accordingly on *match trials* the animal could either make a correct response (‘hit’) or an incorrect response (‘miss’) whereas on *non-match trials* the animal could either make a correct response (‘correct rejection’) or an incorrect response (‘false alarm’). The use of the black circle allowed hit/miss rate to be calculated independently of correct rejection/false-alarm rate; in this way the paradigm is similar to one previously use by Basile and Hampton (Basile and Hampton 2013). A key element of task design in this paradigm for macaque, enabled by the non-match ‘button’ (i.e., black circle), is that miss errors and false-alarm errors may be distinguished independently in which their latencies can be used for estimating the time of familiarity and recollection in macaques. We also varied the degree to which stimuli were either familiar or novel in the session (typically there were up to 80 pairs of novel stimuli that were trial-unique in each session, and six other familiar stimuli, grouped into three pairs, that repeated many times through the session). This allows us to investigate how the image familiarization (repetition of novel sample images) change the relationship between recognition errors and response time.

## Acknowledgements

This work was supported by Wellcome Trust Strategic Award Grant (Ref: WT101092MA) and MRC Project Grant (Ref: MR/K005480/1). We thank the team of expert animal technicians and veterinary staff and anaesthetists for their very high standards of animal care and husbandry throughout.

